# The study of antibody responses to influenza neuraminidase using a lentiviral pseudotype based ELLA

**DOI:** 10.1101/218800

**Authors:** Fabrizio Biuso, George Carnell, Emanuele Montomoli, Nigel Temperton

**Affiliations:** VisMederi S.r.l., Strada del Petriccio e Belriguardo 35, 53100 Siena, Italy.; Viral Pseudotype Unit, University of Kent, Chatham, ME4 4TB, United Kingdom; Department of Molecular and Developmental Medicine, University of Siena, Via Aldo Moro 2, 53100 Siena, Italy.

## Abstract

Influenza pseudotypes represent an alternative to wild type virus for serological assays. To date, pseudotypes (PV) have predominantly been used as surrogates for wild type viruses in microneutralisation assays, where the surface glycoprotein of interest and a reporter gene (such as Luciferase) are used to assess if virus entry into target cells could be inhibited by serum antibodies. The influenza neuraminidase (NA) has the ability to bud and release new virions with or without the contribution of Haemagglutinin (HA). Influenza pseudotypes expressing NA alone, or with HA, were produced to evaluate the antibody response against NA using the enzyme-linked lectin assay (ELLA). The expression of an avian HA with human NAs has enabled the detection of specific antibody reponses against the human circulating subtypes of NA. Within this study a PV-based ELLA assay has been investigated with a pilot panel of sera prepared for an international CONSISE study. Preliminary results have confirmed that the assay is sensitive and could potentially represent a valid alternative to the classical ELLA assay, which requires the employment of reassortant viruses.

## Introduction

Neuraminidase (NA) is the second most abundant glycoprotein on the influenza virus surface (17% of the overall surface) after Hemagglutinin (HA). It is usually expressed at a 1:4 ratio with 40-50 NA tetramers and 160-200 HA trimers on each virion (Knipe and Howley 2007; Air et al., 2012). There are some exceptions, such as the recent pandemic strain (A/California/7/2009 (H1N1), H1N1_/Cal09_) where the ratio may vary (Yang et al. 2012). NA is surrounded by clusters of HA and is responsible for virion release from the host cell surface, promoting the dissociation between HA and sialic acids via its sialidase activity. A direct involvement in influenza infection has been proposed for NA, acting on glycoconjugates present on various lipids and proteins expressed at the cell surface, each having terminal sialic acid residues which could potentially interfere with the receptor binding site of HA. A binding role has also been proposed for NA (Tong et al., 1998; Sung et al., 2010; Yang et al., 2016), especially for some recent H3N2 subtype strains (Lin et al., 2010). Several studies have reported that both inactivated and live attenuated vaccines have the capacity to induce NA-specific antibodies (Hassantoufighi et al., 2010; Couch et al., 2012). These antibodies have the ability to reduce the severity of disease symptoms (Kilbourne 1976) and restrict release and spread of newly formed viruses from infected cells (Compans et al., 1969). This has lead to a growing interest in the development of NA activity detecting assays and neutralisation assays. Several such assays have been developed by utilising different characteristics of the enzymatic reaction to evaluate sialidase activity or antibodies inhibiting it. The thiobarbituric acid (TBA) assay described in 1959 is based on the detection of free sialic acids (Warren 1959). The NA inhibition (NI) assay measures the extent of antibody-mediated interference with enzyme activity (Kilbourne et al. 1968). The NI-TBA assay proposed by the World Health Organization (WHO 2002) and improved by Sandbulte et al. 2009 is considered cumbersome and makes use of hazardous materials (Westgeest et al, 2015). Fluorometric enzyme (FA) and chemiluminescence (CL) assays have been proposed to assess the NI susceptibility of clinical isolates (Wetherall et al., 2003) and have been proposed in the WHO manual (WHO 2011). A QFlu prototype bioluminescence-based NA inhibition assay kit has been used to evaluate the oseltamivir-resistant H1N1pdm09 viruses in clinical specimens, which is not feasible with the other two phenotypic assays without virus culture in cells (Marjuki et al., 2013). Several kits are available on the market but they are expensive and not universally accepted or used. For instance, the National Institute for Medical Research (NIMR) uses a further assay, the fluorescent substrate 2’-(4-methylumbelliferyl)-a-D-N-acetylneuraminic acid (MUNANA) (WHO 2011). A more recent assay, the enzyme-linked lectin assay (ELLA) was developed by Lambre and colleagues (Lambre et al., 1991) and has been further developed and enhanced (Cate et al., 2010; Couch et al., 2012; Couzens et al., 2014). This assay measures the sialidase activity of NA by detecting the terminal galactose that becomes exposed after sialic acid cleavage by NA. A recently study (Eichelberger et al., 2016) made use of the protocol developed by Couzens et al., 2014. It demonstrates that the ELLA assay is robust and sensitive despite issues surrounding the optimal antigen input required. A further study has reported a good intra- and inter-laboratory variability when using a purpose-picked panel of sera, showing that this assay can be standardized but requires further improvements (Eichelberger et al., 2016). Reassortant influenza viruses were used in this study, composed of an H6 HA, internal genes from A/Puerto Rico/8/1938 (H1N1) and N1/N2 NAs from seasonal H1N1 (A/Brisbane/59/2007: H1N1_/BR/07_) or H3N2 (A/Uruguay/716/2007:H3N2_/Ur07_). Unfortunately, as reported by Westgeest et al., 2015, these viruses can be problematic to obtain and handle, making it a challenge for the majority of laboratories to perform this assay. A previous solution to overcome this problem was suggested by Jonges et al., 2010, where wild type H1N1_/Cal09_ X-181 virus was treated with 1% Triton X-100 leading to the dissociation of viral particles, yet maintaining the structure and activity of its NA. Heat treatment was also detailed in this article with satisfactory results. Finally, a simple and innovative solution was recently characterised by Prevato et al., 2015, in a study which was conducted by utilisation of influenza pseudotypes (PV) as a surrogate antigen platform. These PV have already been employed for micronuetralization assays measuring neutralising responses against the HA protein (Temperton et al., 2007; Nefkens et al., 2007). The PV platform avoids the need for reassortant viruses or wild type virus treatment in favor of a faster and safer method of antigen production (Prevato et al., 2015). Preliminary results obtained suggest that the use of influenza PVs for this pseudotype based ELLA (pELLA) could be of increasing interest as already demonstrated for HA neutralising antibodies in the pseudotype based microneutralisation assay (pMN). In this study we have combined the production and assay methods of Prevato et al., 2015 and Couzens et al., 2014, in combination with 8 sera tested and employed in the international CONSISE study for standardization of ELLA (Eichelberger et al., 2016).

## Results & Discussion

As reported by Chen et al., 2007, HA enhances the release of NA from the cell surface and is the rationale for production of influenza PVs expressing both the major surface glycoproteins (HA and NA). In order to emulate reassortant viruses described by Couzens et al., 2014, an expression plasmid bearing the HA gene of A/duck/Memphis/546/74 (H11) was co-transfected with the previously used N1_/Cal09_ and N2_/Tex12_, in combination with p8.91 and pCSFLW plasmids. This enabled the production of H11N1 and H11N2 bearing PV.1µg of NA expression plasmid was used at a 2:1 ratio with HA expression plasmid. The cell medium was replaced 12h post-transfection and harvested 72h post-transfection. Employment of Endofectin™ Lenti lead to increases in NA titres for H11N2 PV (OD_490nm_ increased from 1.91 to 2.36), and similar titres for H11N1 when compared to PEI (OD_490nm_ 3.72 for PEI and 3.45 for Endofectin™ Lenti), shown in Fig.1. In Fig.2, the NA activity results obtained mirror what was found by Chen and colleagues; H11 was able to stabilize and enhance NA activity detected by ELLA.

**Figure 1:**
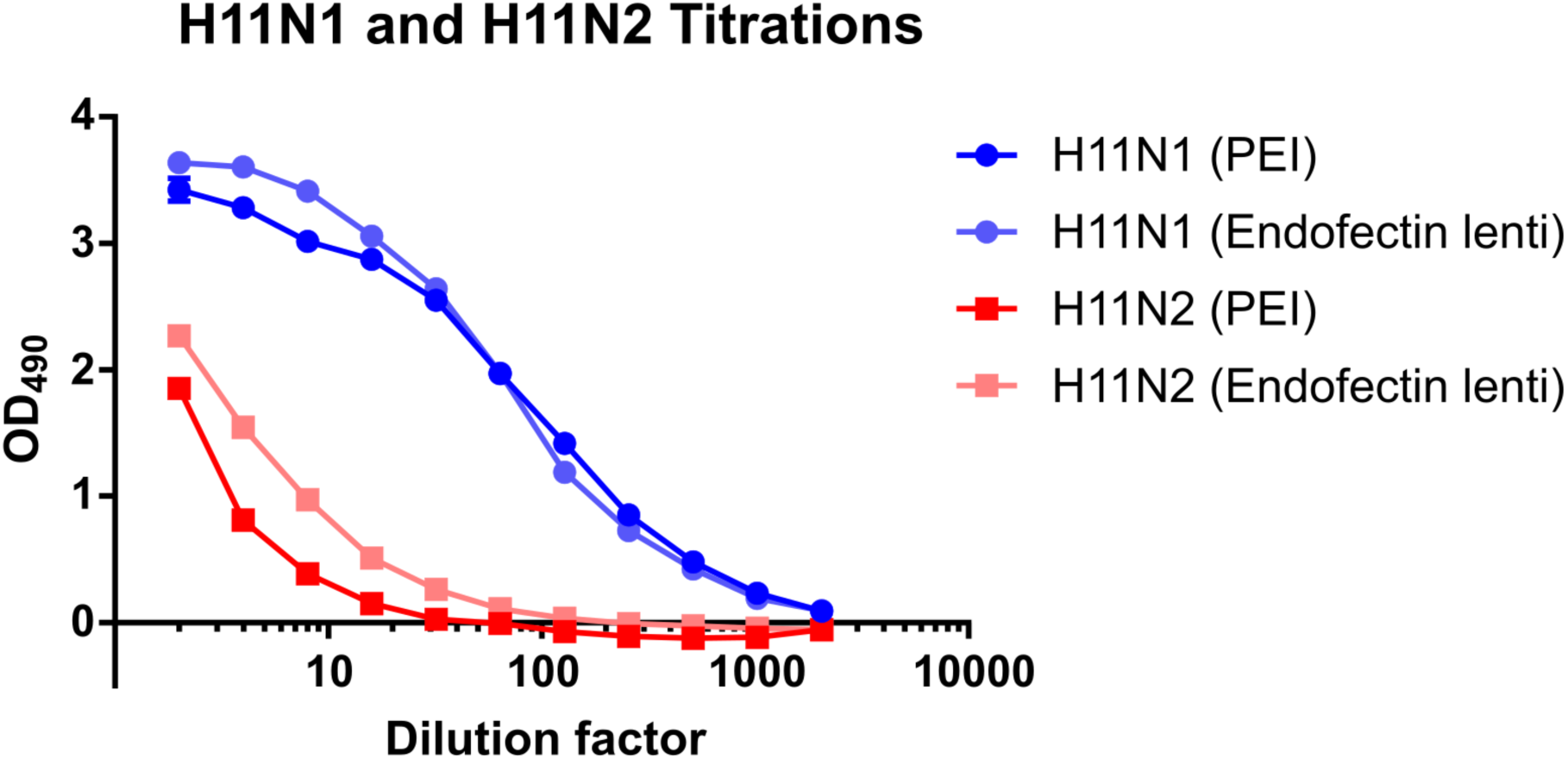
H11N1_/Cal09_ and H11N2_/Tex12_ influenza PV titrations. H11N1_/Cal09_ and H11N2_/Tex12_ PVs obtained by transfecting 0.5µg HA, 1µg NA, 0.5µg gag-pol and 0.75µg of Luc with PEI or Endofectin™ Lenti. On the Y axis is represented the absorbance at 490nm; on the X axis is reported the dilution factor. The use of two alternative reagents was found to result in different yields for H11N2_/Tex12_ PV.

## Influenza NA-bearing PV protocol setup

The positive results obtained from the co-transfection of a non-human HA and human NAs in HEK293T/17 cell lines enabled us to perform a series of pELLA assays in order to evaluate other variables. First, two different concentrations of HRPO were evaluated. The concentration of HRPO reported by Couzens and Eichelberger (Couzens et al., 2014; Eichelberger et al., 2016) was typically 1:1000. Within this study a reduced dilution of HRPO-conjugated lectin (1:500) was used and compared to the standard 1:1000 dilution (Fig. 2). N1_/Cal09_, H11N1_/Cal09_ and H11N2_/Tex12_ PVs exhibited the highest NA activity when a greater amount of HRPO-conjugated lectin was used. This was especially the case for H11N2_/Tex12_ where the higher amount of HRPO was fundamental to perform the assay and analyse the results obtained. At the 1:500 dlution, absorbance values increased. For N1_/Cal09_ and H11N2_/Tex12_PVs the absorbance was almost doubled, while the H11N1_/Cal09_ activity increased less significantly. This demonstrates that the concentration of the conjugate used is not proportional with an increase in NA activity detected. Additionally, differences in OD may be dependent on the different batches of HRPO used.

**Figure 2:**
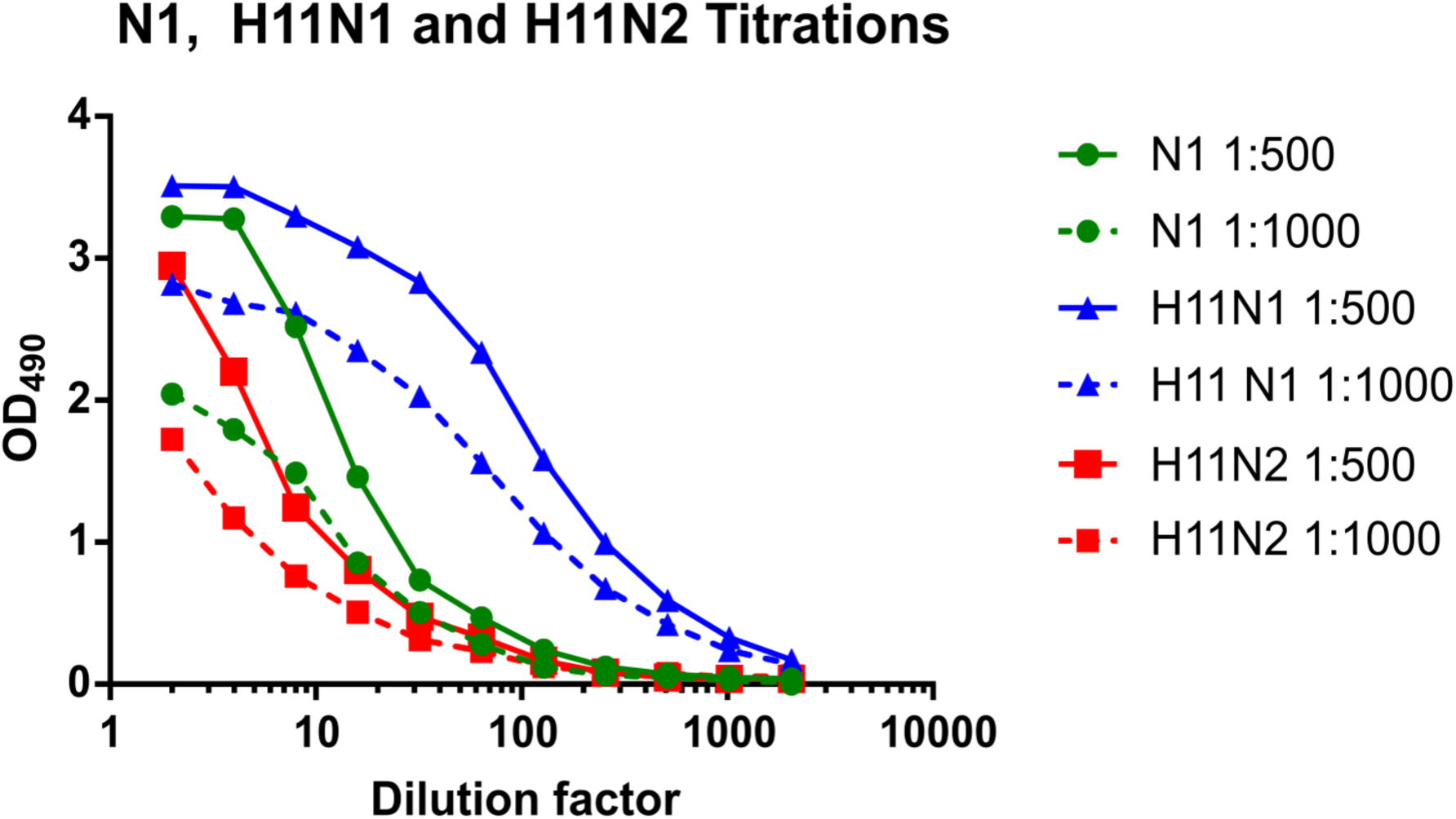
Titration of N1_/Cal09_, H11N1/_Cal09_and H11N2_/Tex12_ influenza PVs. N1_/Cal09_ PVs results obtained by the addition of 1:1000 and 1:500 HRPO-conjugated lectin from Arachisipogea. N1_/Cal09_, H11N1_/Cal09_and H11N2_/Tex12_ influenza PVs are represented as green, blue and red cirlces on lines, resprectively. 1:500 HRPO-conjugated lectin resulted in higher signals for all the three different PVs.

Second, as reported by Eichelberger and colleagues (Eichelberger et al., 2016), the viral input is important when NA inhibition is evaluated, thus two different amounts of antigen have been analyzed in this study; 90-95% of the overall signal, at the start of the linear part of the curve, and OD_490nm_=2.0. This second parameter was chosen according to the input of N1_/Cal09_PVs used by Prevato et al., 2015, corresponding to the start of the linear part of the curve. As reported by Eichelberger et al., 2016, absorbance values ranging from <1.7 to 4 did not show any significant difference in geometric mean titer (GMT) or percent of geometric coefficient of variation (%GCV) when measuring antibody titres by ELLA.

## PVs-based ELLA (pELLA) neutralisation assay

The protocol adopted for the evaluation of NA inhibition was described previously (Couzens et al., 2014). Deviations from this method include the use of 1:500 as well as 1:1000 dilutions of HRPO-conjugate. 8 serum samples (S001-S008) which derive from a CONSISE study (Eicheberger et al., 2016) were tested in duplicate by two independent operators. The inhibition curves generated were highly reproducible for all PV tested (Fig 3). The trends of serum NA inhibition are generally conserved between the three different influenza PV strains, with high responses against H11N1_/Cal09_ corresponding with N1_/Cal09_and H11N2_/Tex12_. The end-point titre was calculated as the reciprocal of the highest dilution at which the serum is able to induce 50% inhibition of the maximum titre (VC) obtained (Table 2).

**Figure 3:**
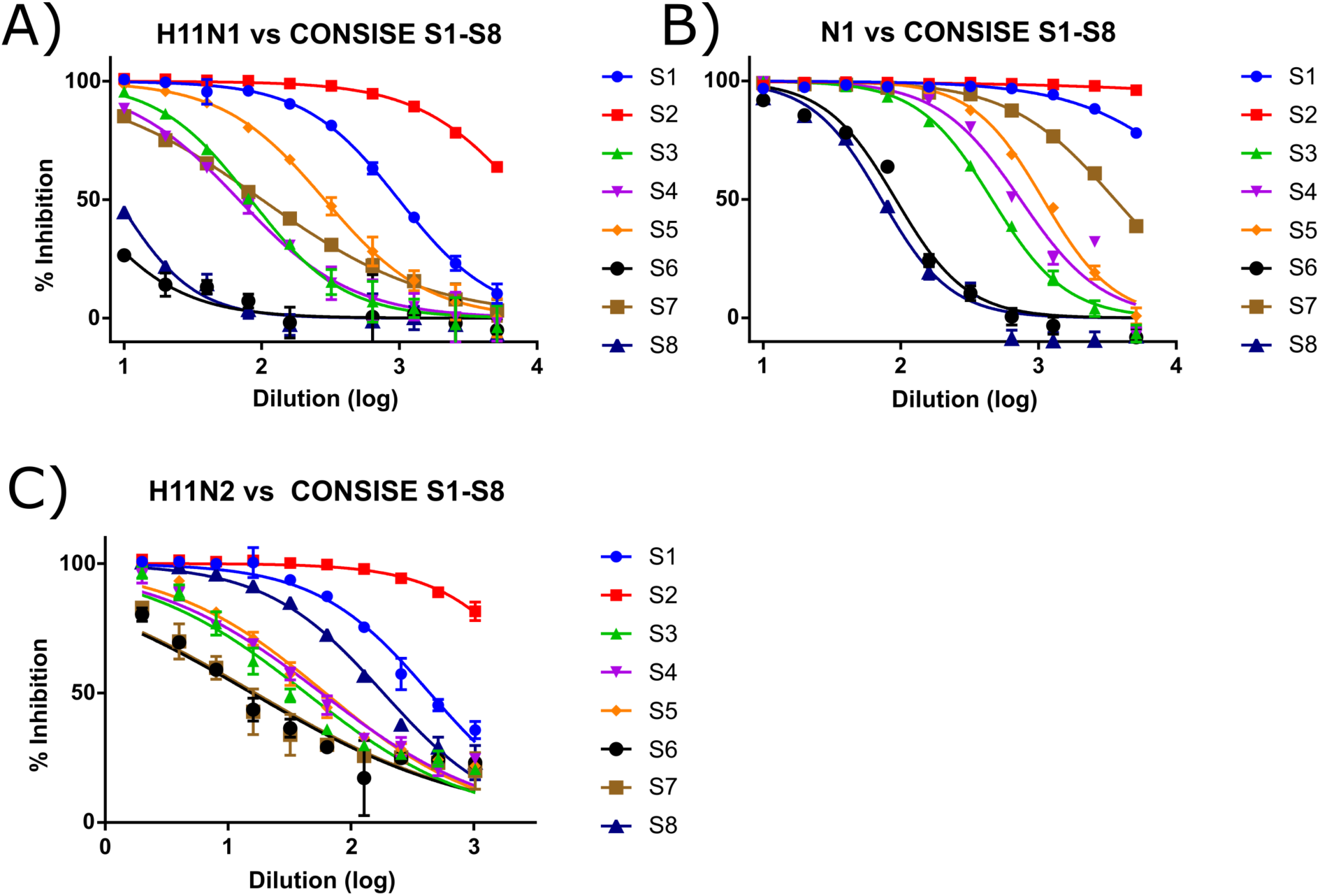
Inhibition curves generated by pELLA, conducted with different influenza NA-bearing PVs: S1-S8 sera analyzed with N1_/Cal09_ PV (A), with H11N1_/Cal09_ PV (B) and with H11N2_/Tex12_ PV (C). Each antigen was tested in duplicate.

**Table 2:** End-point titres obtained by pELLA neutralisation assay of 8 serum samples. H11N1_/Cal09_ and H11N2_/Tex12_ PVs used.

| SERUM<br>ID | N1 |  | H11N1 |  | H11N2 |  |
| --- | --- | --- | --- | --- | --- | --- |
|  | Titre |  | Titre |  | Titre |  |
| S001 | >5120 | >5120 | 640 | 640 | 2560 | 1280 |
| S002 | >5120 | >5120 | >5120 | >5120 | >5120 | >5120 |
| S003 | 640 | 640 | 80 | 80 | 160 | 160 |
| S004 | 1280 | 640 | 160 | 160 | 320 | 320 |
| S005 | 1280 | 1280 | 160 | 160 | 320 | 320 |
| S006 | 160 | 160 | <10 | <10 | 80 | 80 |
| S007 | 5120 | 5120 | 40 | 40 | 40 | 80 |
| S008 | 80 | 80 | <10 | <10 | 1280 | 1280 |

S002 was able to strongly inhibit NA activity of both NA antigens used for the pELLA assay, especially against N1_/Cal09_ PV. Overall, inhibition titres are significantly increased when sera are used against N1_/Cal09_ PV as opposed to H11N1_/Cal09_ PV, as shown in Table 2. These results are interesting as the PVs produced by Prevato et al., 2015 expressed only NA on their surface. These data show that while the expression of HA may enhance PV NA activity titres, it interferes with inhibition titres. This may be through steric hindrance which could hinder antibodies binding to the NA stem or masked regions. This effect has been reported by HA-stalk targeting antibodies (Rajendran et al. 2017).

As reported in Table 1 the sera tested have different origins. S001 was obtained from human purified immunoglobulin selected to have high HAI antibody titers against recent circulating influenza strains and the pELLA results confirm this, showing a minimum titre of 640 when tested against H11N1_/Cal09_ PV. S002 derives from bovine plasma collected from cows that are transgenic for human immunoglobulins, vaccinated with 2012/2013 trivalent inactivated influenza and show the highest titre (>5120) against all three PV used. The pooled human serum of sample S003 showed different antibody titres against the three PV tested. The S004-S008 samples were collected from adult volunteers that had received the seasonal 2009/2010 influenza vaccination. The fluctuation in results is most likely due to each patient’s immune history and response to the 2009/2010 vaccine.

**Table 1:** Sera used for the pELLA assay. This table was adapted from Eichelberger et al., 2016.

| SAMPLE ID | CHARACTERISTICS |
| --- | --- |
| S001 | Human IVIG (Baxter Biomedical Research); purified immunoglobulin from donors selected in 2013 to have high HAI antibody titers against recent circulating influenza strains |
| S002 | Transgenic bovine plasma (Sanford); plasma was collected from cows that are transgenic for human immunoglobulins and had been vaccinated with 2012/2013 trivalent inactivated influenza vaccine. |
| S003 | Pooled human sera (Lonza) |
| S004-S008 | Human serum (Accelovance); each sample was collected from an adult volunteer post seasonal 2009/2010 influenza vaccination. |

Interestingly, these sera show higher antibody titres against H11N2_/Tex12_, which contains the NA from a strain isolated in 2012, than against the H11N1_/Cal09_ and N1_/Cal09_PV. Since the H1N1_/Cal09_ virus originated from a triple reassortant virus with a swine NA of Eurasian avian-like origin (Tang et al., 2010), it may explain why sera display higher antibody titres against N2_/Tex12_. There is a distinct difference in antibody titres against N1_/Cal09_ and H11N1_/Cal09_ PVs. The N1_/Cal09_ PV lacks any HA on its surface, leaving the NA exposed. Thus, NAs can be bound by antibodies which may not have been able to do so in the presence of HA. Expressing NA by itself may also decrease the stability of the glycoprotein, as has been suggested in another study regarding HA-bearing PV (Oh et al. 2009), a lower number of NA expressed on PVs could enhance the ability of sera to block and impede sialidase activity. However, the co-expression of H11 with N1 could have i) improved the number of NA particles released (Chen et al., 2007) ii) stabilized the HA-NA expression maintaining the optimal ratio (Chen et al., 2007) iii) blocked the antibodies directed against certain regions of NA which would be unable to bind in the presence of HA. In regards to H11N2_/Tex12_ or N2_/Tex12_ titres, it could be hypothesized that the lower titres obtained from the titration could be due to a lower presence of particles released, a lower expression of NA on the PV surface or suboptimal conditions for the activity of the N2 enzyme.The recent findings of Gao et al., 2016 suggest that a more acidic environment stabilizes the N2 protein, increasing the reliability of downstream results.N2_/Tex12_ could also simply have a lower activity which would be reflected in our results.

An attempt to align results obtained in this study with those produced by the CONSISE group (Eichelberger et al., 2016) was carried out. It is of importance to note that there are a number of differences between the two sets of antigens used within these studies and thus data generated by linear regression could not be compared between studies. The GMT of the results obtained from this study was compared with those obtained from the CONSISE study (Table 3). Reassortant H6N1_/Br07_ and H6N2_/Ur07_viruses (HA gene from A/turkey/Massachusetts/3740/1965 (H5N2)) were used as antigens in the ELLA assay. Thus, different NAs were used, both from 2007, while the influenza PVs produced express N1 from H1N1_/Cal09_ and N2 from H3N2_/Tex12_.

**Table 3:** N1_/Cal09_, H11N1_/Cal09_ and H11N2_/Tex12_ PVs (violet) pELLA results compared to H6N1_/Br07_ and H6N2_/Ur07_ (HA gene from A/turkey/Massachusetts/3740/1965 (H6N2), N1 and N2 antigens of seasonal influenza virus H1N1_/Br07_ and H3N2_/UR07_ respectively: orange) ELLA results.

| SERUM ID | N1(GMT) | H11N1(GMT) | H6N1(GMT) | H11N2(GMT) | H6N2(GMT) |
| --- | --- | --- | --- | --- | --- |
|  | Titre | Titre | Titre | Titre | Titre |
| S001 | >5120 | 640 | 817 | 1810 | 680 |
| S002 | >5120 | 5120 | 121 | >5120 | 3293 |
| S003 | 640 | 80 | 23 | 160 | 31 |
| S004 | 905 | 160 | 70 | 320 | 42 |
| S005 | 1280 | 160 | 110 | 320 | 46 |
| S006 | 160 | 5 | 9 | 80 | 9 |
| S007 | 5120 | 40 | 22 | 57 | 34 |
| S008 | 80 | 5 | 11 | 1280 | 147 |

Some differences are seen in the results obtained with different NA strains. S001 from N1_/Cal09_ has a very high antibody titre in respect to H11N1_/Cal09_ and H6N1_/Br07_, which are within a one-log dilution difference. H11N2_Tex12_ and H6N2_/Ur07_ differ by more than one-log dilution but both show high antibody titres. S002 antibody titres are more than four-fold higher against N1_/Cal09_ and H11N1_/Cal09_ than H6N1_/Br07_, meaning that the immunoglobulins contained in transgenic bovine plasma react very differently between these strains, most likely as the vaccine used is matched to the strain used in the pELLA and not the original study. Serum sample S002 shows high titres against both N2s. The GMT antibody titres against N1_/Cal09_ PVby S003 exhibited a very high end-point when compared to those generated against H11N1_/Cal09_ and H6N1_/Br07_, while there is less than a two-fold difference in dilution between antibody titres for the latter two antigens. The results for H11N2_Tex12_ and H6N2_/Ur07_ underline that there are at least two-fold dilutions between the two different antigens used. S004 shows a similar trend to S003. S005 shows high antibody titres against N1_/Cal09_ but similar titres between H11N1_/Cal09_ and H6N1_/Br07_, while there is at least a three-fold difference in antibody titre between H11N2_/Tex12_ and H6N2_/Ur07_. S006 is shown to have a very low affinity for all the viruses tested and, while its antibody titre against N1_/Cal09_ is five-fold higher than against H11N1_/Cal09_ and H6N1_/Br07_, it shows the same antibody titre against the latter two. S007 antibody titres are similar between H11N1_/Cal09_ and H6N1_/Br07_ as well as between H11N2_/Tex12_ and H6N2_/Ur07_. S008 exhibits the lowest antibody titre againstN1_/Ca09_, but it is still greater than three-fold dilutions higher than against H6N1_/Br07_, which is in-line with H11N1_/Cal09_. However, S008 shows a very high antibody titre againstH11N2_/Tex12_, which is around three-fold higher than for H6N2_/Ur07_.

It is important to underline that while the results obtained by pELLA originate from sera tested in duplicate, the GMTs used as comparison were produced by 23 different labs. To better estimate the magnitude of differences between the results obtained by the present study and from the already published work, a further comparison will have to be made. As shown in Table 3 and already confirmed by the end point titres, the N1_/Cal09_ PV are more readily inhibited in all cases when compared to H6N1_/Br07_ results with the same sera. However, H11N1_/Cal09_ PV titres are within the same range as H6N1_/Br07_ results in 7/8 cases. Only S002 tested against H11N1_/Cal09_ PV shows values significantly different when compared to the ones obtained by using the reassortant H6N1_/Br07_ virus. However, antibody titres generated against H11N2_/Tex12_were only similar to H6N2_/Ur07_ in 2/8 cases, with the former showing higher inhibition. While it is difficult to compare the results generated between each study, they reaffirm that pELLA could be used to detect antibodies against NA on its own, to generate data free from any HA directed antibody interference.

For the H1N1_/Cal09_ strain, the NA and matrix genome segments originate from Eurasian avian-like H1N1 swine influenza viruses, while the other six segments are derived from North American triple-reassortant, H1N1/H1N2/H3N2 swine influenza viruses (Tang et al., 2010). The amino acid sequence alignement conducted using the SIM Alignment Tool (ExPASy web.expasy.org/sim/) shows that N1_/Br07_ and N1_/Cal09_ share 81.5% identity (Fig. 4). Nevertheless, the results evaluated by calculating the end-point titres show similar trends.

**Figure 4:**
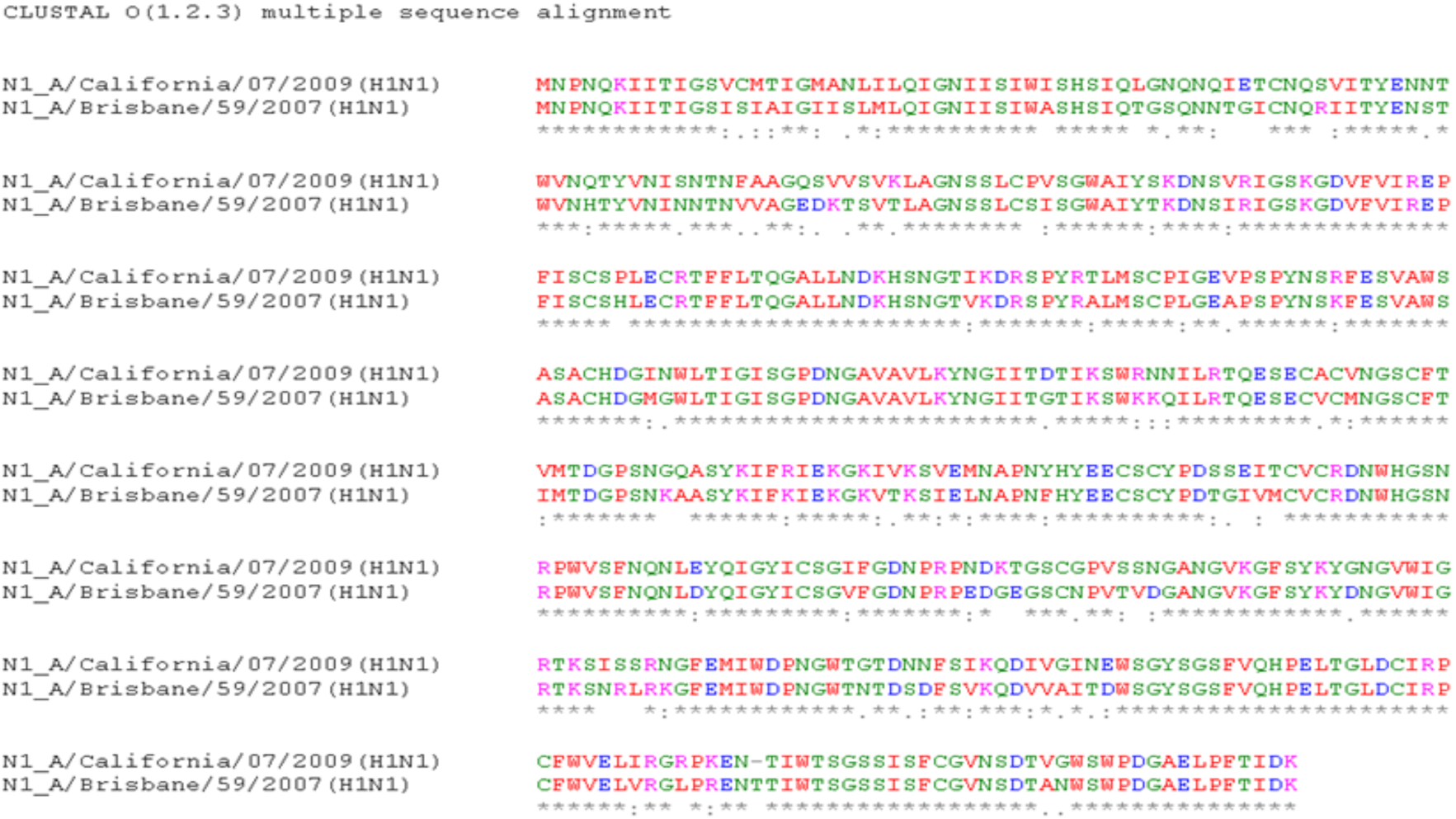
N1 proteins (H1N1_/Cal09_ and H1N1_/Br07_ alignment by using Clustal Omega (http://www.ebi.ac.uk/Tools/msa/clustalo/).

The same alignment analysis was conducted for H3N2_/Ur07_ and H3N2_/Tex12_ (Fig. 5). Unlike N1, these two sequences show a higher identity (97.7%, evaluated by SIM Alignment Tool - ExPASy). Reassortant viruses H6N1_/Br07_ and H6N2_/Ur07_ were chosen for the international study as they were antigenically similar to the vaccine composition strains for the 2009-2010 influenza season. It is likely that the sera tested against N2 are more reactive against H11N2_/Tex12_ than H6N2_/Ur07_ despite the similarity in amino acid sequence.

**Figure 5:**
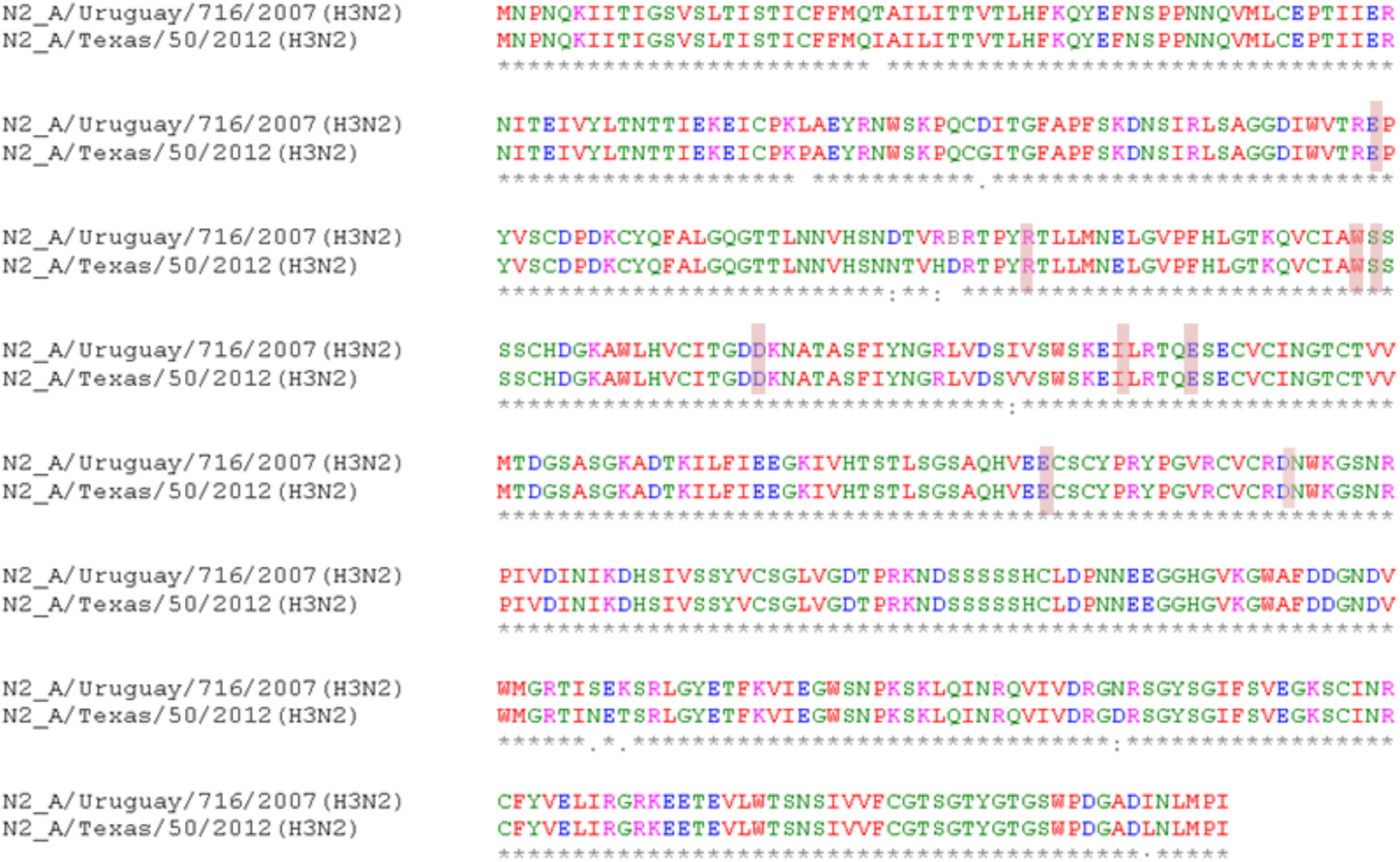
N2 proteins (H3N2_/Ur07_ and H3N2_/Tex12_) alignment by using Clustal Omega (http://www.ebi.ac.uk/Tools/msa/clustalo/). The active site residues are highlighted in pink

An alignment between the NA sequence of H3N2_/Ur07_, H3N2_/Pth09_ and H3N2_/Tex12_ using Clustal Omega shows that there are only 11 amino acid mutations between all three strains. The H3N2_/Ur07_ NA differs from that of H3N2_/Pth09_ by 4 amino acids, while there are 7 amino acid differences between it and H3N2_/Tex12_. The hypothesis that H3N2_/Pth09_ is more similar to H3N2_/Ur07_than H3N2_/Tex12_ is confirmed by the percent identity matrix shown in Table 4. The HAI results reported by the WHO (http://www.who.int/influenza/vaccines/200909_Recommendation) show an increasing proportion of post-infection ferret antisera antigenic and genetically distinguishable from the previous vaccine viruses used (H1N1_/Br07_ and H3N2_/Ur07_) in favour of a close relation to H3N2_/Pth09_ and H3N2_/HK09_ reference viruses.This may explain why the sera obtained from patient’s post-seasonal 2009/2010 influenza vaccination (S004-S008) show higher antibody titres against N2_/Tex12_ than N2_/Ur07_, even though these HAI results are related to HA evolution.

**Table 4:** Percent identity matrix between H3N2_/Ur07_, H3N2_/Pth09_ and H3N2_/Tex12_ strains

| % identity | N2<sub>/Tex12</sub> | N2<sub>/Ur07</sub> | N2<sub>/Pth09</sub> |
| --- | --- | --- | --- |
| N2<sub>/Tex12</sub> | 100 | 97.87 | 98.51 |
| N2<sub>/Ur07</sub> | 97.87 | 100 | 98.93 |
| N2<sub>/Pth09</sub> | 98.51 | 98.93 | 100 |

To evaluate this hypothesis, a deeper analysis of the amino acid differences between the H3N2_/Ur07_ and H3N2_/Tex12_ viruses was performed. Few mutations lie close to the critical and highly conserved residues among all the influenza subtypes, apart from the D147N substitution where an aspartic acid residue has mutated to an asparagine from H3N2_/Ur07_ to H3N2_/Tex12_. Also, a negative residue has changed into a polar uncharged residue among the NAs. This conserved residue, Asn146 is glycosylated and located in close proximity to the active site of NA (Shtyrya et al., 2009). The glycans are reported to have a regulatory function in relation to NA enzymatic activity (Shtyrya et al., 2009). Thus, it is possible to hypothesize that the D147N substitution in close proximity to this site, changing the charge of the side chain, may have somehow decreased the avidity of antibodies for H3N2_/Ur07_ NA in favour of H3N2_/Tex12_.

**Figure 6:**
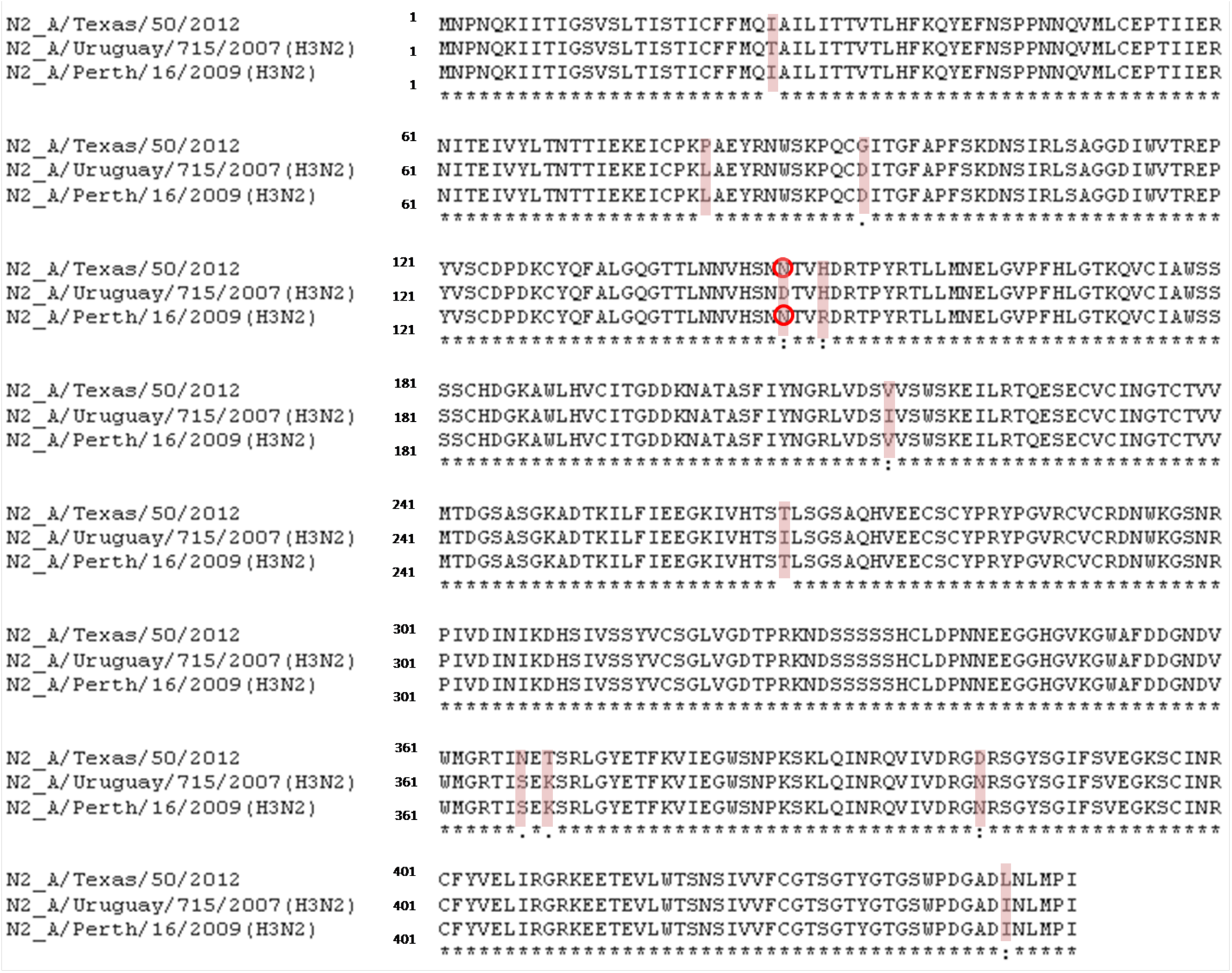
N2 proteins (H3N2_/Ur07_, H3N2_/Pth09_and H3N2_/Tex12_ aligned by using Clustal Omega (http://www.ebi.ac.uk/Tools/msa/clustalo/). The 11 mutations occurring between the strains are highlighted in pink. The substitution D147N is shown as red empty circles.

In conclusion, we can speculate that i) patients S004-S008 could have been infected naturally with new circulating strains increasing the avidity of sera for N2_/Tex12_ but more realistically ii) there is high cross-reaction coupled with iii) greater instability of N2-bearing PVs leading to these higher antibody titres. However, several further analyses should be performed and a comparison between reassortant viruses matched with influenza PV should be carried out using the same NA.

## Conclusion

The aim of this study was to evaluate the utility of influenza PVs that bear NA, in order to be used as surrogate viruses for serological assays. Despite multiple different methodologies that have been proposed throughout the years for the detection of sialidase activity, there is a growing interest for ELLA, which was proposed in 1991 and then further adapted (Lambre et al., 1991; Couzens et al., 2014). The first study using PVs in ELLA was described recently (Prevato et al., 2015). However, in this present study it is possible to better understand the behaviour of influenza NA-bearing PVs in the pELLA assay and some of the critical variables involved. Some success in the production of higher titre NA only PVs in comparison to the first report has been achieved (Prevato et al., 2015). Results obtained from the CONSISE study (Eichelberger et al., 2016) aimed to evaluate the putative standardization of this method. The results obtained are very encouraging since by following the the protocol described by Couzens et al., 2014, it was possible to asses a wide range of anti-NA antibody responses, resulting in more robust outputs than the those obtained by Prevato et al., 2015. From this study it emerges that NA itself is able to form PVs without the need of HA, but that HA increases PV release and leads to overall higher NA titres (Chen et al., 2007; Nayak et al., 2009; Rossman and Lamb 2011). While we have not confirmed similarities between influenza WT virus and PV morphology by scanning electron microscope (SEM) or transmission electron microscopy (TEM), previous studies have confirmed this (Chen et al., 2007; Wu et al., 2010; Yang et al., 2014; Prevato et al., 2015) and titrations have yielded positive results. The assay, as described in Couzens et al., 2014, is shown to be sensitive and robust and this study confirms this. There was no difference in 50% cut-off titre when HRPO was used at at 1:1000 or 1:500 dilutions. On the other hand, as discussed by Eichelberger et al., 2016, the viral input used to perform the assay is fundamental. In the aforementioned study it was suggested that the viral input should be 90-95% of the maximum OD_490nm_ signal reached in the titration, which should correspond to the start of the linear part of the curve. In this study, we compared this input (OD_490nm_ of ~3) to the one used by Prevato et al., 2015, where the 90-95% of the maximum OD_490nm_ corresponded to 2. The results (not shown) were conserved between each condition tested (4-fold increase in antibody titres), but were not comparable in terms of actual antibody titre (IC_50_ cutoff). The NA inhibition titres obtained from an OD_490nm_ of 2 are higher, meaning that sera inhibit NA with a greater magnitude. Thus, the relationship between viral input and corresponding IC_50_ or end-point titre values must be evaluated. In conclusion, the existing ELLA protocol is suitable for the use of PV in order to generate NA inhibition antibody titres. However, a further comparison of ELLA/pELLA by using the same set of sera against the same NAs but from different sources (reassortant virus and PV) should be performed. It would be of significant interest and value to assay the same set of sera against different NAs of the same group (i.e. group 1) to evaluate putative cross-reacting immune responses.

## Materials and methods

### Plasmids, cells

HEK293T/17 cells and phCMV1-H11 (A/duck/Memphis/546/1974 (H11N9)) were kindly provided by Dr DavideCorti, Institute for Research in Biomedicine, Switzerland. N1_/Cal09_and N2 (A/Texas/50/2012 (H3N2), H3N2_/Tex12_) were synthesised (WT sequence) and subcloned into plasmid expression vector pI.18 (Cox et al. 2002) by Genscript (U.S.A). HIV-1 derived packaging plasmid pCMVΔR8.91 (p8.91) was obtained from Dr Yasu Takeushi, University College London, originating from the laboratory of Dr Didier Trono (Naldini et al. 1996; Zufferey et al. 1997). Plasmid pCSGW originated from (Demaison et al. 2002), the GFP gene was replaced by that encoding firefly luciferase, forming the plasmid pCSFLW by Novartis Vaccines, Italy.

### Production of PV

Production of lentiviral PV was carried out by co-transfection of HEK293T/17 cells using branched polyethylenimine (PEI) or Endofectin Lenti™ transfection reagents. Briefly, a DNA mix of 0.5 µg of phCMV1-H11, 0.5 µg of p8.91, 0.75 µg of pCSFLW and either 1 µg of N1_/Cal09_ or N2_/Tex12_ was mixed in 200µl Opti-MEM™ and incubated for 20min with PEI (17.5µl of 1mg/ml PEI per well of a 6-well plate) or 25 min with Endofectin Lenti™ (3 µl/µg of DNA) per well of a 6-well plate. This mix was transferred to 60-90% confluent HEK293T/17 cells. Cell culture medium (DMEM +10% FBS +1% Penicillin/Streptomycin) was changed after overnight incubation at +37°C 5% CO_2_ in a humidified incubator. Cells were then left for a further 48h and supernatants harvested 72h post transfection, and sterile filtered through a 0.45µm filter (Merck Millipore).

### Sera

The sera used within this study were kindly provided by Dr Maryna Eichelberger and were part of an international study carried out by the CONSISE group (Eichelberger et al., 2016). 8 sera (S001-S008) were pre-treated for 1h at 56°C and used at a 1:10 starting dilution in the ELLA inhibition assay.

### ELLA Titration assay

Titration of NA activity was performed as described previously (Couzens et all., 2014; Prevato et al. 2015). Briefly, a starting volume of 240µl of PV containing supernatant was serially diluted (1:2) across a 96-well clear plate in sample diluent, resulting in wells with 120µl of sample diluent and a dilution gradient of PV. 50µl of this was transferred to 96-well maxisorp plates previously coated in fetuin, in duplicate. 90-95% of the maximum OD_490nm_ signal, corresponding to the start of the linear part of the curve, was used as viral input to run the ELLA neutralisation assay.

### ELLA inhibition assay

The characteristics of each serum sample tested (Barr et al., 2010) are reported in Table 1 (Eichelberger et al., 2016). Briefly, 24µl of serum was added to 216µl of sample diluent giving a starting dilution of 1:10 in a 96-well clear plate. 120µl of sample diluent was aliquoted into all other wells on the plate and serial 1:2 dilutions were carried out, giving a dilution gradient of 1:10 to 1:5120 in volumes of 120µl. 50µl of the contents of each well were then transferred to a 96-well maxisorp plate previously coated in fetuin. This was performed in duplicate, giving two data points per well of the serum dilution series. 50µl of antigen source (PV) was then added to wells in row 1 to 11 in order to give the desired OD_490nm_ based on previous titration results. The 12^th^ row was filled with 50µl of sample diluent as a background control. Plates were then incubated overnight (o/n) at +37°C 5% CO_2_ and then washed 6x with wash buffer (PBS containing Tween^®^ 20 at 0.05%) before the addition of 100µl of conjugate diluent containing the peanut agglutinin PNA-HRPO (1:500 or 1:1000 dilution). The plate was then incubated for 2h at room temperature (RT) in the dark. Subsequently, the plate was washed 3x with wash buffer and 100µl of substrate (OPD dissolved in citrate buffer) was added to each well. 100µl of stop solution was added immediately before results were read with a Tecan Sunrise microplate reader (10min incubation). Results were read at OD_490nm_.

## Acknowledgements

We declare that there is no conflict of interest in the production of this article. We would like to thank Maryna Eichelberger and Laura Couzens for providing sera and reassortant viruses for use in ELLA. H6N1 and H6N2 reassortant viruses were prepared using plasmids that were a kind gift to Maryna Eichelberger from St Jude Children’s Research Hospital, Memphis, Tennessee, USA. We would also like to thank Vadim Sumbayev and Isabel Gonzalez Silva (Medway School of Pharmacy, UK) for their assistance with the spectrophotometeric analysis.

## Author contributions

FB and GC designed and carried out the experiments under direction from NT and EM. All authors prepared and reviewed the manuscript prior to submission.

